# Highly controllable co-delivery of siRNA and doxorubicin via cationic niosomes for synergistic anticancer effects

**DOI:** 10.1101/2025.09.04.674162

**Authors:** Ping Song, Tianqiang Liu, Kurt V. Gothelf, Jørgen Kjems

**Affiliations:** Center for DNA Nanotechnology (CDNA) and Interdisciplinary Nanoscience Center (iNANO), Center for RNA Therapeutics towards Metabolic Diseases (RNA-META), Aarhus University, Gustav Wieds Vej 14, DK-8000 Aarhus C, Denmark; Department of Molecular Biology, University of Aarhus, DK-8000 Aarhus, Denmark; Department of Chemistry, Aarhus University, Langelandsgade 140, DK-8000 Aarhus C, Denmark; Hainan Institute of Northwest A&F University, Sanya, Hainan Province 572024, China

**Keywords:** Co-delivery, Doxorubicin, siRNA, Niosome, Oligodeoxynucleotides

## Abstract

The combination of anticancer drugs and siRNA has emerged as a promising strategy for cancer therapy due to their synergistic effects. However, a major challenge is developing a co-delivery system with high controllability capable of simultaneously encapsulating both the chemical drug and siRNA. In this study, we developed a simple and highly controllable co-delivery system for delivering Doxorubicin (Dox) and siRNA. Co-encapsulation was achieved by intercalating Dox into duplex GC-rich oligodeoxynucleotides, (CGA)_7_/(TCG)_7_ (termed CGA-Dox), followed by mixing with siRNA and subsequent condensation using a cationic noisome (SPN) through electrostatic interactions. SPN/siRNA/Dox exhibited efficient cellular uptake and high gene silencing efficiency in both H1299 and MDA-MB-231 cells. Furthermore, co-delivery of Dox and siRNA targeting ribonucleotide reductase subunit M2 (RRM2) using this system significantly enhanced cellular apoptosis and antiproliferative effect in MDA-MB-231 cells compared to Dox alone. These results indicate that our co-delivery system constructed via Dox-DNA intercalation and electrostatic complexation, represents a promising strategy for synergistic anticancer therapy.

## 1 Introduction

In recent years, most cancer therapies relying on a single chemotherapeutic drug have proven unsatisfactory due to the development of drug resistance, metastasis, mutations and activation of alternative cancer progression pathways induced by adaptive responses during the treatment ^1–3^. Combination therapy has become a widespread approach to enhance anti-tumor sensitivity, improve therapeutic efficiency and minimize side effects, particularly when combining chemotherapeutic agents with gene therapy.

To overcome multidrug resistance and achieve improved therapeutic efficiency, small interfering RNA (siRNA) has been employed to block the translational of genes involved in multidrug resistance or the anti-apoptotic cellular defense^4^–6. Ribonucleotide reductase subunit M2 (RRM2) has long been recognized as an important therapeutic target for diseases depending on DNA replication, including cancer. RRM2 is overexpressed in various cancer cells and plays a crucial role in the development of tumor invasion and malignancy by promoting the DNA synthesis, repair and cell proliferation. Overexpression of RRM2 is associated with drug resistance and inhibition of Ribonucleotide reductase (RR) has shown potential anticancer effect^7,8^. For DNA-damaging anticancer drugs, suppressing RRM2 has shown significant overturn of drug resistance ^9^. Several delivery materials have been developed for the co-delivery of siRNAs and anticancer drugs, with various systems based on inorganic nanoparticles^10^, lipoplexes ^11^, polyplexes ^12,13^, dendrimers ^14^, liposomes and micelles^15–17^. However, preparation of these multifunctional delivery systems is generally complicated, involving multiple steps and difficulty in controlling the physiochemical properties. This often results in heterogeneous drug-loading capacity, payload leaking, siRNA degradation during preparation, and compromised reproducibility^18^. Therefore, there is a strong need to develop a simple system capable of encapsulating siRNA together with anticancer drugs in a highly controlled manner.

DNA-based platforms composed of GC-rich oligodeoxynucleotides have been applied for efficient loading of Dox. For example, Kim *et al*. developed an aptamer with a GC-rich duplex tail for targeted Dox delivery, achieving high drug-loading capacity through intercalation of Dox into CG base pairs^19,20^. The negatively charged Dox-intercalated DNA oligonucleotides (CGA-Dox) could be efficiently encapsulated with cationic carriers via electrostatic interactions, enabling one-step co-encapsulation of siRNA and CGA-Dox through the same mechanism. Non-ionic surfactant based vesicles, known as “niosomes”, have been reported as an alternative to liposomes due to their higher stability and loading capacity^21,22^. Moreover, cationoic noisomes are relatively non-toxic and exhibited efficient siRNA delivery properties^23,24^. In this study, we prepared a sorbitan monooleate (Span) - based cationic niosome (SPN), composed of the non-ionic surfactant Span80, the cationic lipid 1,2-dioleoyl-3-trimethylammonium-propane (DOTAP) and D-α-Tocopheryl Polyethylene Glycol 1000 Succinate (TPGS), to simultaneously encapsulate CGA/Dox and siRNA via electrostatic interactions. The in vitro performance of SPN/siRNA/Dox in terms of characterization, cellular uptake, transfection and gene-silencing efficiency was evaluated in the highly invasive breast cancer cell line MDA-MB-231 and the non–small-cell lung cancer cell line H1299. We further applied SPN/siRRM2/Dox to deliver Dox and silence RRM2 in MDA-MB-231. The cytotoxicity and apoptosis induction of the co-delivery system were subsequently evaluated to confirm the synergetic anti-cancer efficiency.

## 2 Materials and methods

### 2.1 Materials

Doxorubicin hydrochloride, D-α-tocopheryl polyethylene glycol-1000 succinate (TPGS) and Span 80 were purchased from Sigma-Aldrich. 1,2-dioleoyl-3-trimethylammonium-propane (DOTAP) was obtained from Avanti Polar Lipids (Alabaster, AL). siRNAs against GFP (siGFP): sense, 5’- GACGUAAACGGCCACAAGUTC-3’, antisense, 5’-ACUUGUGGCCGUUUACGUCGC-3’; siRNA against RRM2 (siRRM2): sense, 5’-GAUUUAGCCAAGAAGUUCAGA-3′, antisense, 5′- UGAACUUCUUGGCUAAAUCGC-3’ and Cy5-labeled siGFP were obtained from Ribotask (Odense, Denmark). The scrambled negative control siRNA (siNC) was provided by Genepharm (Shanghai, China), sense, 5’-UUCUCCGAACGUGUCACGUTT-3’, antisense, 5’- ACGUGACACGUUCGGAGAATT-3’. (CGA)7/(TCG)7 oligodeoxynucleotides (sequence: 5’- CGACGACGACGACGACGACGA-3’, complementary sequence: 5’- TCGTCGTCGTCGTCGTCGTCG-3’) were purchased from IDT (Integrated DNA Technologies, Inc.)

### 2.2 Cell culture

The human breast cancer cell line MDA-MB-231 was maintained in RPMI medium supplemented with 10% fetal bovine serum, 1% penicillin-streptomycin at 37°C in 5% CO_2_ and 100% humidity. H1299 cells stably expressing green fluorescent protein (H1299-GFP) were kindly provided by Dr. Anne Chauchereau, (CNRS, Villejuif, France), and cultured in growth medium (RPMI) with 500 μg/ml Geneticin (Gibco, Life Technologies).

### 2.3 Synthesis of SPN and SPN/siRNA/Dox complex

The cationic niosome was prepared by the so called ethanol injection method according to our previous report^24,25^. Briefly, DOTAP, Span 80 and TPGS were dissolved in ethanol at concentrations of 50, 100 and 100 mg/ml, respectively. The lipid mixture was combined at the molar ratio of 50:45:5 and injected quickly into distilled water with vigorous stirring to generate empty vesicles (SPN). To develop SPN/siRNA/Dox, Doxorubicin (Dox) and CGA oligodeoxynucleotides were dissolved in PBS at the concentration of 100 μM and 20 μM respectively and mixed at different molar ratios and incubate for 30 min at room temperature to produce CGA-Dox complexes. Then, siRNA (20 μM) and the prepared CGA-Dox were mixed at a molar ratio of 4:1, 2:1 or 1:1, followed by complexation with SPN at various weight ratios and incubation for 30 min at room temperature. Freshly prepared SPN/siRNA/Dox complexes were further utilized for further characterization and cell experiments.

### 2.4 Characterization of SPN and SPN/siRNA/Dox complex

The size and zeta potential were measured by a Zetasizer Nano ZS (Malvern Instruments, Malvern, UK) at 25 °C. The morphologies of SPN, SPN/siRNA, SPN/siRNA/CGA and SPN/siRNA/Dox were monitored by Transmission Electron Microscope (TEM, FEI Tecnai G2 Spirit). Samples were prepared by adding 3 μl sample suspension to a carbon filmed Cu grid and stained with uranyl formate.

The encapsulation of Dox was characterized by a fluorimeter. Briefly, the fluorescence spectrum of SPN/siRNA/Dox was acquired at excitation of 480 nm and emission from 520 nm to 750 nm using a Horiba Jobin Y von Fluoremax-3 fluorimeter. The encapsulation efficiency of siRNA was measured by RiboGreen reagent (Invitrogen, Copenhagen) according to our previous reports^26^. Briefly, 50 uL of SPN/siRNA/CGA solution with 50 uL of a diluted RiboGreen solution (1:200 in TE buffer) was mixed in a 96-well plate (Nunc, Roskilde, Denmark). After incubating for 5 min, the fluorescent intensity was measured using a FLUOstar OPTIMA (BMG, Labtechnologies) at excitation and emission wavelengths of 480 and 520 nm, respectively.

### 2.5 Cellular uptake

MDA-MB-231 cells were seeded in 8-well cell culture chambers (SARSTED, Germany) at 2.5×10^4^ cells/well. After overnight incubation, the medium was changed with fresh medium containing SPN/Cy5-siRNA/Dox, SPN/Cy5-siRNA and Lipofectamine 2000/Cy5-siRNA. The final concentration was 50 nM for siRNA and 0.25 μM for Dox, respectively. After incubation for 8 h, cells were fixed with 4% paraformaldehyde for 15 min at room temperature, washed three times with PBS and mounted with ProLong® Gold Antifade Mountant containing DAPI stain (Cat. P36941, Life Technologies). Cellular uptake was imaged using confocal microscope LSM710

(Zeiss, Germany). To quantify cellular uptake, MDA-MB-231 cells were seeded in 24 well-plate (5×10^4^ cells/well) and incubated overnight. The different complexes described above were added to the cells. After 8 hours’ incubation, the cells were detached with 0.05% Trypsin-EDTA (Gibco, Invitrogen) and Cy5 or Dox fluorescence intensity was quantified by flow cytometry (Becton Dickenson FACSCalibur).

### 2.6 Knockdown of GFP

H1299-GFP cells were seeded in a 24-well plate at 5×10^4^ cells/well and incubated overnight. SPN/siRNA, SPN/siRNA/Dox, and Lipofect/siRNA were added to the cells at final concentrations of 50 nM for siRNA and 0.25 μM for Dox, respectively. After overnight incubation, the medium was changed with fresh medium and incubate for another 6 h. The cells were then visualized and imaged at FITC channel by fluorescent microscope (OLYMPUX IX71) and silencing efficiency of GFP was further quantified by flow cytometry (Becton Dickenson FACSCalibur).

### 2.7 Knockdown of RRM2

MDA-MB-231 were seeded in 24-well cell plates at 5×10^4^ cells/well and incubated overnight. The cells were administrered with SPN/siRRM2/Dox, SPN/siNC/Dox, SPN/siRRM2, SPN/siNC, Lipofect/siRRM2 and Lipofect/siNC. The final concentration was 50 nM for siRNA and 0.25 μM for Dox, respectively. After overnight incubation, the medium was changed with fresh medium and incubated for another 6 h. Total RNA was extracted with Trizol reagent (Invitrogen) according to the manufacturer’s protocol. cDNA was then synthesized by using RevertAid RT Reverse Transcription Kit (Thermo Scientific) and used as template for quantitative PCR using LightCycler® 480 SYBR Green(Roche) on a LightCycler® 480 Real-Time PCR system (Roche).

Relative gene expression level was normalized to the housekeeping gene glyceraldehyde-3-phosphate dehydrogenase (GAPDH), and data represented an average of duplicates per experiment. Primers used for qPCR were shown in Supplementary Material (Table S1).

### 2.8 Cytotoxicity analysis

The *in vitro* cytotoxicity of SPN/siRRM2/Dox against MDA-MB-231 cells were measured by Alamar Blue assay (Molecular Probes, Life Technologies) according to the manufacturer’s protocol. MDA-MB-231 cells were seeded in 96-well plate (1×10^4^ cells/well) and incubate overnight. The cells were then treated with SPN/siRRM2/Dox, SPN/siRRM2, SPN/siNC/Dox, SPN/siNC and free Dox. The final concentration of siRNA was 50 nM and Dox 0.25 μM respectively. After 24 hours’ incubation, cells were rinsed with PBS and replaced with an Alamar Blue reagent (10% in growth media) for 2 h at 37 °C. Fluorescence intensity of the medium was measured using a plate reader (FLUOstar OPTIMA, Moritex Bioscience) at λex of 540 nm and λem of 590 nm (n=5).

### 2.9 Cell apoptosis analysis

Annexin V-FICT/PI apoptosis detection kit (Sigma-Aldrich) was used to quantify apoptosis according to the manufacture’s recommendation. MDA-MB-231 Cells were seeded in 24-well plate and transfected with SPN/siRNA, SPN/siRNA/Dox complex and free Dox at the final concentration of 50 nM for siRNA and 0.125 μM for Dox. After 48 hours, cells were trypsinised, washed with PBS, centrifuged, resuspended in 300 μl of 1X Annexin V-binding buffer supplemented with 2.5 μl Annexin V-FITC and 2.5 μl PI (Propidium iodide) (50 μg/ml). Samples were incubated at 4 °C for 30 min with protection from light, and detection of viable cells (Annexin V-FITC negative, PI negative), apoptotic cells (Annexin V-FITC positive, PI negative), and necrotic cells (Annexin V-FITC positive, PI positive) was performed by using a FACSCalibur (Becton Dickinson).

### 2.10 Statistical analysis

Data are presented as means ± standard deviation (SD). Statistical significance was analyzed using one-way ANOVA with a LSD post-hoc test. Differences were considered significant when p < 0.05.

## 3 Results

### 3.1 Characterization CGA-Dox complex

CGA-Dox complex was prepared by incubating Dox with the GC-rich oligodeoxynucleotide duplex in PBS at room temperature. To confirm Dox intercalation, the fluorescence spectra of free Dox and CGA-Dox were recorded at an excitation wavelength of 480 nm and an emission range of 520 - 750 nm. Free Dox exhibited a strong fluorescent emission peak around 600 nm, whereas the fluorescence of Dox in CGA-Dox was quenched (Figure S1A), likely due to π-π stacking interactions between Dox and CG bases resulting from the close molecular proximity^27,28^. The maximum loading ratio of (CGA)_7_/(TCG)_7_ to Dox was achieved at 0.125, indicating that one mole of oligonucleotides could intercalate approximately 8 moles of Dox.

### 3.1 Characterization SPN and SPN/siRNA/Dox complex

Empty SPN vesicles were synthesized by injecting a lipid mixture of DOTAP, Span-80 and TPGS into aqueous solution, where they self-assembled into cationic niosomes through hydrophobic interactions. SPN exhibited well-defined properties with an average size of 151.4 ± 6.82 nm and polydispersity index (PDI) of 0.19 ± 0.02 (Figure 1A). Subsequently, siRNA and CGA-Dox were mixed and condensed with SPN to generate SPN/siRNA/Dox complexes. Encapsulation efficiency of siRNA and CGA-Dox was determined using RiboGreen assay, which measures the fluorescent intensity of unbound nucleic acids. As shown in Figure S1, complete binding of siRNA and CGA-Dox was observed when the SPN-to-(siRNA + CGA-Dox) weight ratio reached 15, indicating full condensation of siRNA and CGA-Dox by SPN. The result aligns with our previous study, and thus, we will utilize this ratio for subsequent experiments due to its sufficient gene-silencing efficiency and biocompatibility^24^. To confirm Dox encapsulation, fluorescence spectra of SPN/siRNA/Dox were compared with those of free Dox at the same concentrations. As shown in Figure 1D, SPN/siRNA/Dox displayed nearly complete quenching at all the three concentrations (0.125 μM, 0.25 μM and 0.5 μM), indicating that Dox intercalation into the CGA oligo was maintained during SPN complexation.

**Figure 1.**
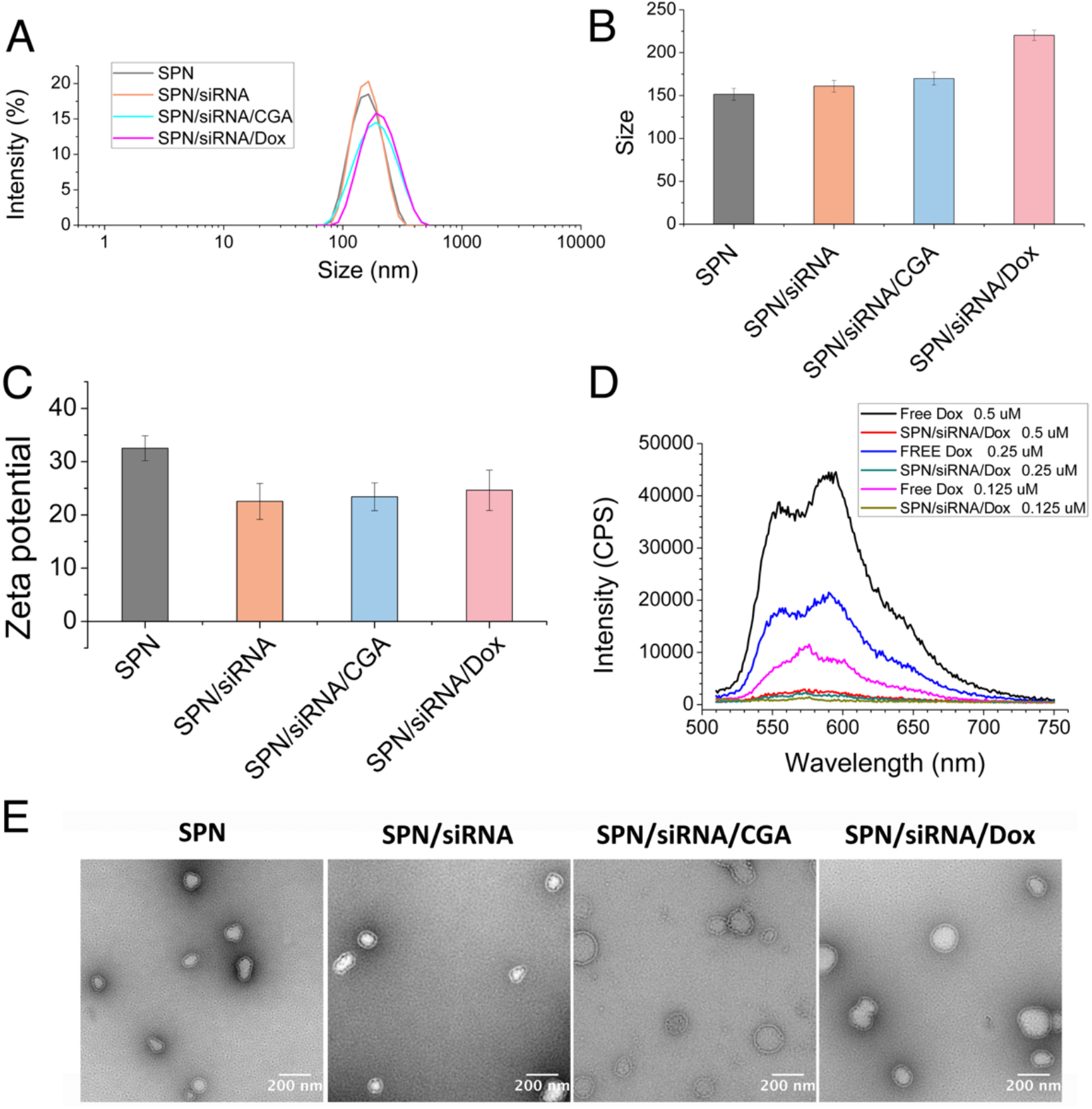
Characterization of SPN, SPN/siRNA, SPN/siRNA/CGA and SPN/siRNA/Dox. (A) Hydrodynamic size distribution of indicated formulations by DLS. (B) Quantification of size from DLS measurement (n=3). (C) Zeta potential measurements of the indicated formulations. (D) Fluorescence emission spectra of SPN/siRNA/Dox compared to free Dox at Dox concentration of 0.5 μM, 0.25 μM and 0.125 μM, respectively. (E) TEM images of SPN, SPN/siRNA, SPN/siRNA/CGA and SPN/siRNA/Dox. Scale bar, 200 nm.

The physiochemical properties of SPN-based delivery systems were further characterized after loading of Dox and siRNA. The sizes of the SPN based delivery systems slightly increased after the loading of siRNA, CGA, or CGA-Dox, respectively (Figure 1A, B). In addition, the surface charge of SPN (32.5±2.34 mV) decreased due to the encapsulation of negatively charged siRNA and CGA molecules. In general, after complexing with siRNA and CGA/Dox (SPN/siRNA/CGA-Dox), the particles remain slightly positively charged (24.6±5.95 mV) and with a hydrodynamic size of 220.2± 5.95 nm with a PDI of 0.24±0.02 (Figure 1C). Furthermore, the TEM images confirmed the well-dispersed particles with good uniformity and liposome-like structure (Figure 1E).

### 3.3 In vitro cellular uptake

Cellular uptake of SPN/siRNA/Dox in MDA-MB-231 cells was evaluated using confocal microscopy and quantified by flow cytometry. As shown in Figure 2A, cells treated with SPN/Cy5-siRNA/Dox displayed significantly stronger intracellular Cy5 signals than those treated with Lipofect/Cy5-siRNA, indicating enhanced cellular siRNA uptake mediated by SPN. Comparable Cy5 fluorescence intensities were observed between SPN/Cy5-siRNA and SPN/Cy5-siRNA/Dox treatment, suggesting that introduction of CGA/Dox did not impair siRNA uptake. Additionally, Dox was localized in the nucleus of cells treated with SPN/siRNA/Dox. Flow cytometry quantification was consistent with confocal microscopy results: SPN-based systems demonstrated higher Cy5 intensity compared with Lipofect/Cy5-siRNA (Figure 2B, C). Moreover, SPN/Cy5-siRNA/Dox-treated cells exhibited higher nuclear Dox accumulation compared with cells treated with free Dox (Figure 2D, E).

**Figure 2.**
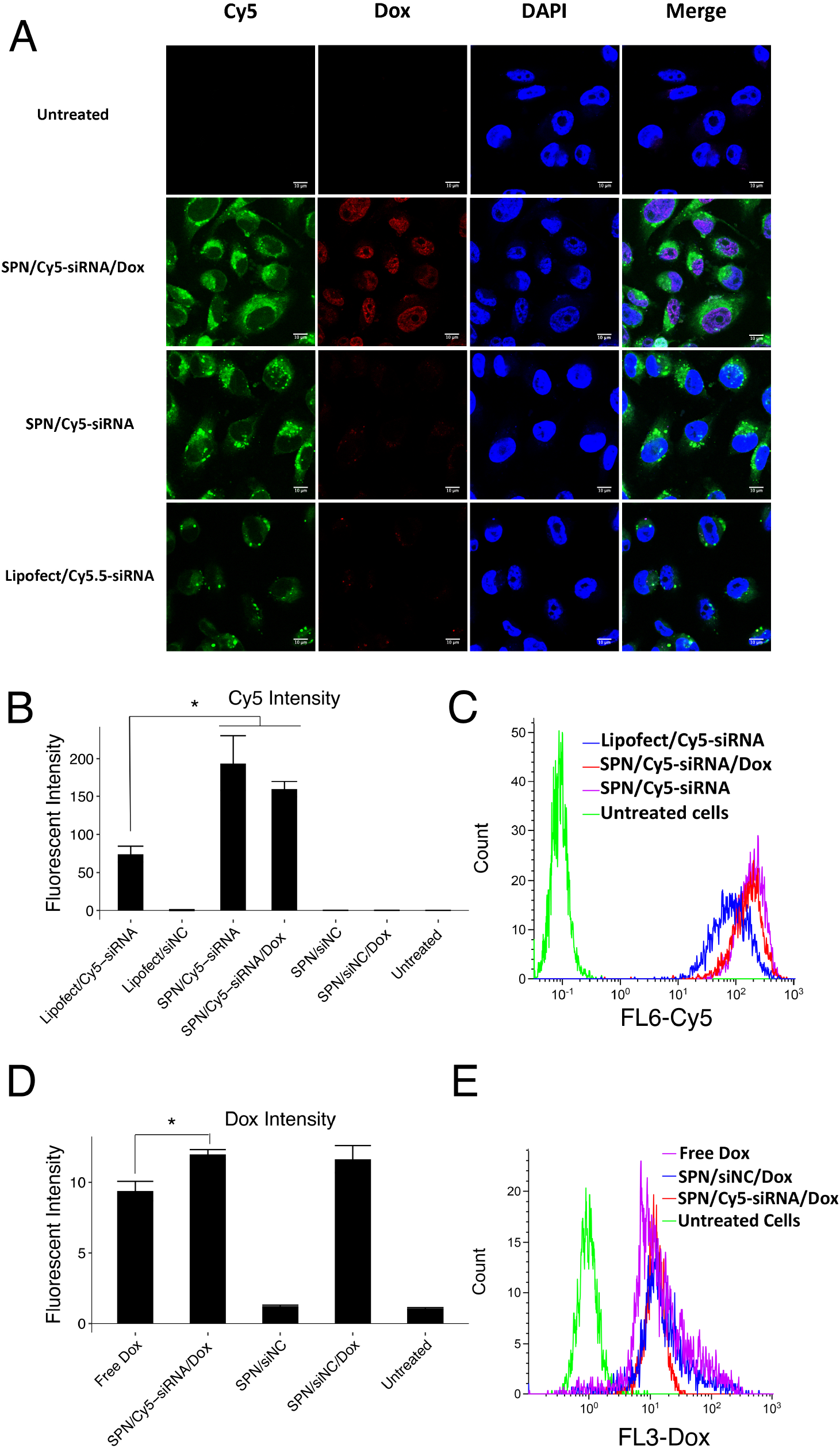
Uptake of SPN/siRNA/Dox by MDA-MB-231 cells. (A) Representative confocal images of untreated cells and cells transfected with SPN/Cy5-siRNA/Dox, SPN/Cy5-siRNA or Lipofect/Cy5-siRNA for 8 hours. Nucleus were stained by DAPI (Blue), Cy5-siRNA and Dox were indicated in Green and Red, respectively. Scale bar: 10 μm (B) Quantification of Cy5-siRNA uptake in cells treated with indicated formulations by flow cytometry (n=3, * *p < 0*.*05*). (C) Representative histogram indicating the Cy5-siRNA uptake. (D) Quantification of Dox uptake by flow cytometry (n=3,* *p < 0*.*05*). (E) Representative histogram indicating the uptake of Dox.

### 3.4 Gene silencing efficacy

To assess the gene-silencing efficacy of the co-delivery system, we first encapsulated siRNA against green fluorescent protein (GFP) (siGFP) and evaluated GFP knockdown in H1299-GFP cells. A significant decrease in GFP signals was observed in cells treated with SPN/siGFP and SPN/siGFP/Dox compared with negative controls (SPN/siNC and SPN/siNC/Dox) (Figure3A). Silencing efficiency was quantified by flow cytometry: SPN/siGFP/Dox achieved approximately 74% GFP knockdown, comparable to the commercial transfection reagent Lipofectamine 2000 (Figure 3B,C).

**Figure 3.**
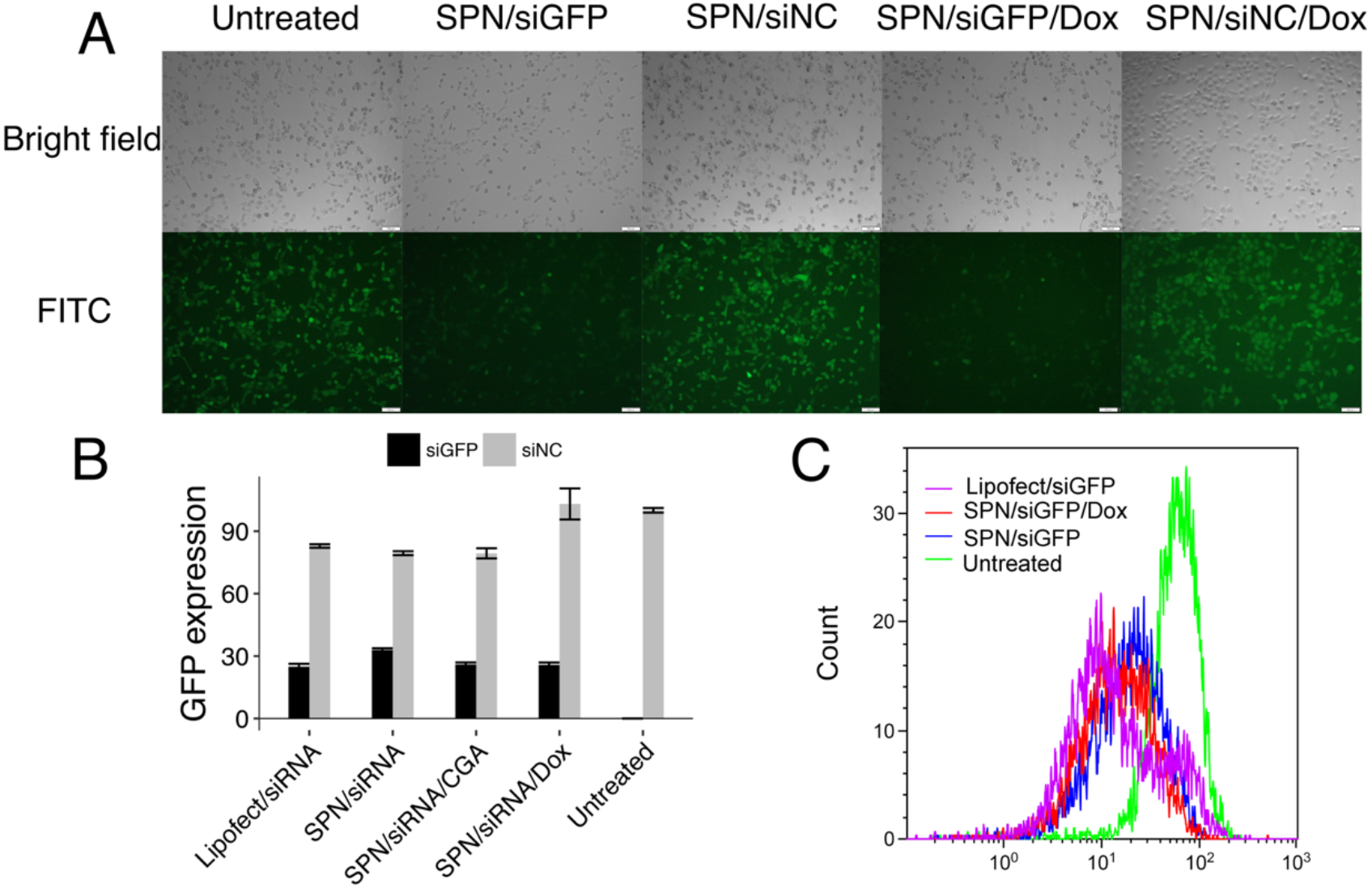
Knockdown of GFP in GFP expressing H1299 cells. (A) Representative fluorescent images of H1299 cells after 24-h treatment of indicated formulations. Scale bar 100 μm; (B) Quantification of relative GFP expression by flow cytometry, Mean values were normalized to percent control (n=3). (C) Representative histogram of flow cytometry of GFP fluorescent intensity in H1299 cells after treatment with SPN/siGFP/Dox, SPN/siGFP and Lipofectamine 2000, respectively. Untreated cells (medium only) served as the negative control, representing normal GFP levels.

RRM2 is one of the two subunits of ribonucleotide reductase that regulates DNA synthesis and cell proliferation^9^. Thus, RRM2 knockdown combined with DNA-damaging chemotherapeutics such as Dox may produce a synergistic effect. To confirm the capability of silencing RRM2, SPN/siRRM2/Dox, SPN/siRRM2 and their negative controls (SPN/siNC/Dox and SPN/siNC) were tested in MDA-MB-231 cells. As shown in Figure 4, SPN/siRRM2/Dox achieved approximately 84% knockdown, significantly higher than Lipofect/siRRM2 (approximately 74%). Furthermore, silencing efficiency was similar between SPN/siRRM2/Dox and SPN/siRRM2, indicating that CGA-Dox encapsulation did not hinder siRNA-mediated gene silencing. Untreated cells (medium only) were used as the negative control to represent normal mRNA levels, ensuring that any reduction in expression observed in treated groups was due to specific knockdown rather than nonspecific toxicity of the SPN-based systems.

**Figure 4.**
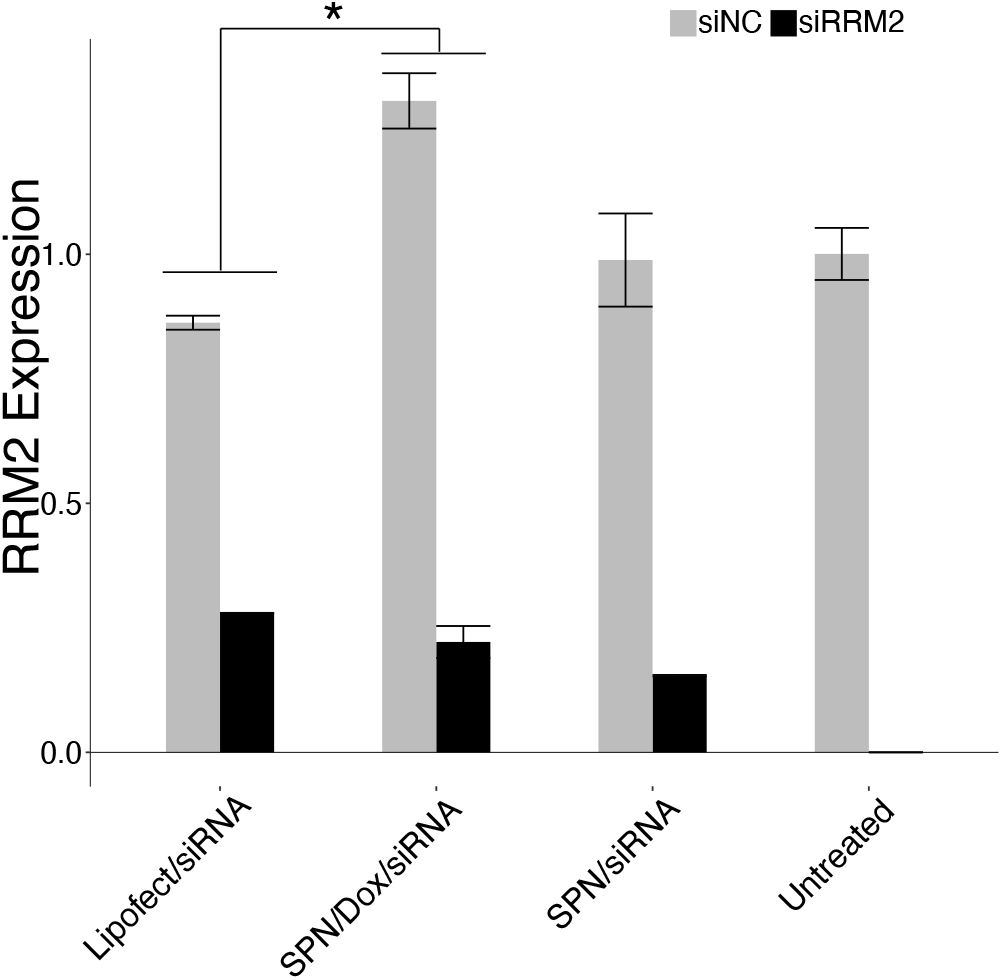
Knockdown of RRM2 in MDA-MB-231 cells. Relative RRM2 mRNA expression in MDA-MB-231 cells was quantified by qPCR after 24-h treatment with the indicated formulations., Relative RRM2 mRNA expression was normalized to GAPDH. Untreated cells (medium only) served as the negative control, representing normal RRM2 mRNA levels. (mean ± SD, n = 3, \**p < 0*.*05*)

### 3.5 Cytotoxicity analysis

To determine whether RRM2 knockdown enhances the chemotherapeutic effect of Dox, cytotoxicity can be increased after knockdown of RRM2 by the co-delivery system. Cytotoxicity after 24 hours was measured by AlamarBlue essay in MDA-MB-231. As revealed in Figure 5, SPN/siRRM2/Dox exhibited significantly higher cytotoxicity on MDA-MB-231 cells (79.0 %) than free Dox (26.6%). In addition, SPN/siRRM2/Dox demonstrated significantly higher toxicity compared with SPN/siNC/Dox (*p < 0*.*001*), which confirmed the enhanced anticancer effect from synergic effect of RRM2 silencing and Dox delivery. To be noted, SPN/siNC did not show much toxicity to the cells, indicating the good biocompatibility of SPN.

**Figure 5.**
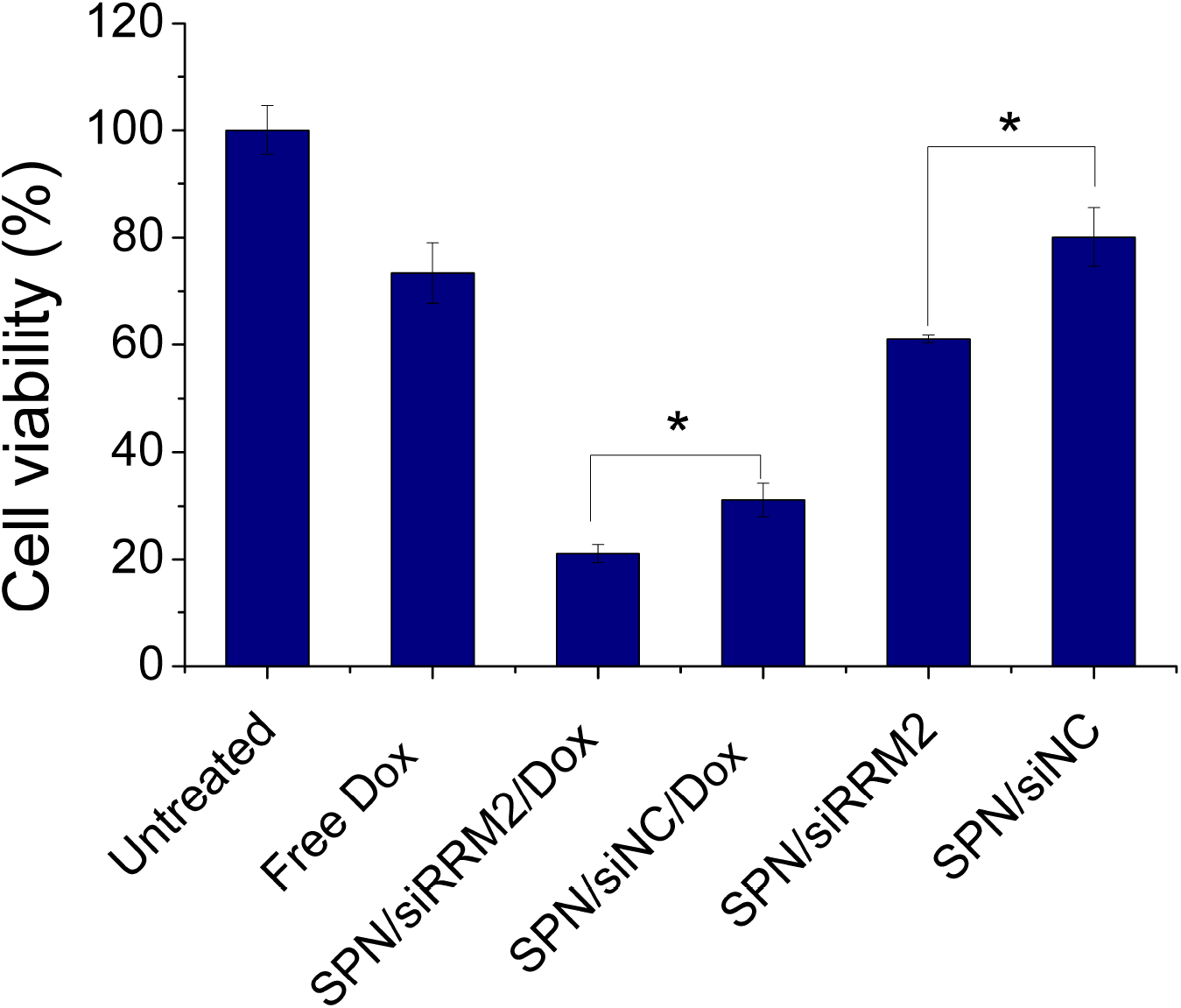
Cell viability of MDA-MB-231 cells treated with SPN/siRNA, SPN/siRNA/Dox, SPN/siNC, SPN/siNC/Dox and free Dox for 24 hours, in which siRNA represented siRRM2. Data was expressed as mean ± SD (n=5). Significance: *\* p < 0*.*05*.

### 3.6 Cell apoptosis

To investigate the pro-apoptotic effect of SPN/siRRM2/Dox, Annexin V/PI staining was performed. Annexin V binds to phosphatidylserine (PS), which is normally located on the inner layer of cell membrane but is translocated to the outer membrane during apoptosis and exposed to be easily detectable. Propidium iodide (PI) penetrates only damaged or dead cells to stain DNA, enabling differentiation between viable and nonviable cells^29^. Cells negative for both Annexin V and PI (Annexin V^−^/PI^−^) are considered viable; Annexin V+/PI^−^ negative cells are early apoptotic cells; and Annexin V^+^/PI^+^ cells are late apoptotic or necrotic. As show in Figure 6, SPN/siRRM2/Dox treatment significantly reduced viable cells (66.1 ± 3.5%) and increased apoptotic cells (17.5 ± 0.2 %) compared with SPN/siNC/Dox (viable: 72.7 ± 3.1%; apoptotic: 13.1 ± 1.7 %). SPN/siRRM2 alone slightly decreased viable cells (84.5 ± 1.9%) relative to SPN/siNC (90.1% ± 1.6%), with apoptotic cells rising from 4.2 ± 1.1% (SPN/siNC) to 7.9% ± 2.2% (SPN/siRRM2). Necrotic cells (late apoptotic) increased significantly in both Dox-encapsulated systems and the free Dox groups, while only 3.3 % of apoptotic cells were observed with free Dox, likely due to activation of resistance pathways preventing apoptosis.^33^ Importantly, SPN/siNC did not significantly reduce the cell viability compared with untreated cells, suggesting that SPN itself is biocompatible and does not adversely affect cell growth. Overall, these results indicate a synergistic effect of SPN/siRRM2/Dox in inducing apoptosis in cancer cells.

**Figure 6.**
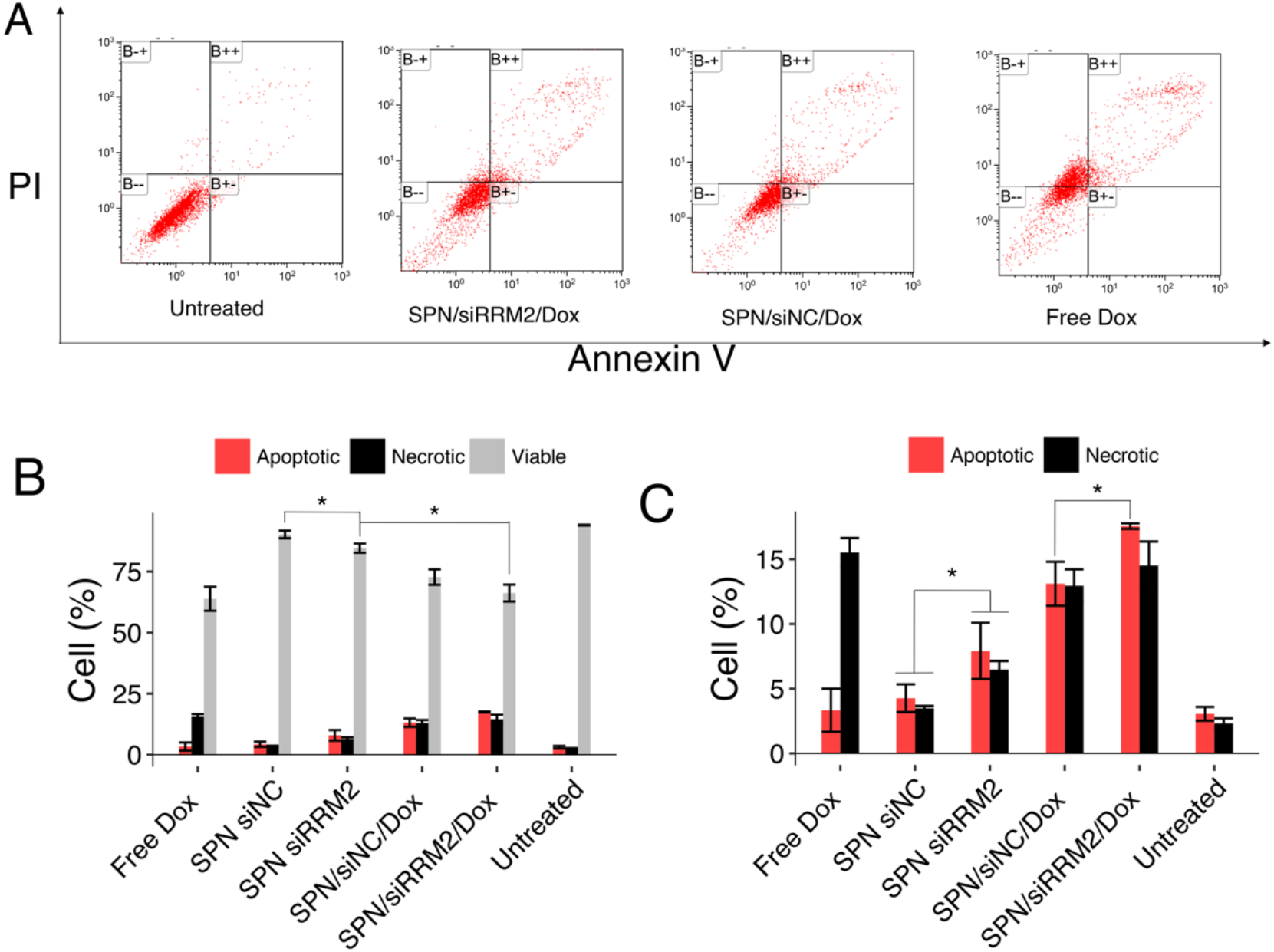
Apoptosis analysis of MDA-MB-231 cells after treatment with indicated formulations. (A) Histogram of flow cytometry analysis of apoptosis, cells were stained with Annexin V-FITC /PI, (B--), (B+-) and (B++) represent viable, apoptotic and necrotic cells, respectively; (B) Quantification of the percentage of viable, apoptotic and necrotic cells from flow cytometry analysis; (C) Amplify the percentage of apoptotic and necrotic cells. Data was expressed as mean ± SD (n=3). Significance: * *p < 0*.*05*.

## Discussion

Combination strategies involving chemotherapeutics and RNAi-based gene therapy have been investigated for many years. Currently, most existing co-delivery systems usually employ sequential encapsulation of chemotherapeutics in vesicles followed by gene encapsulation. This strategy makes it challenging to maintain reliable physicochemical properties and reproducibility. In addition, such systems may suffer from drug leakage and siRNA degradation during storage, which further limits the clinical translation. Therefore, great challenges remain in achieving flexible and well-controlled co-loading of drug and gene. In this study, we utilized a 21-base-pair DNA duplex (CGA)7/(TCG), which demonstrated high capacity for Dox intercalation with good stability and preserved anti-tumor effect. Our co-delivery system utilizes the ionic nature of both CGA-Dox and siRNA, enabling one-step encapsulation by cationic vesicles through electrostatic interactions.

In cellular uptake studies, SPN/siRNA/Dox exhibited high fluorescent signals for both siRNA and Dox in MDA-MB-231, confirming the efficient co-delivery and intracellular release. CGA/Dox complexes are reported to be highly stable with negligible leakage in the blood circulation^31,32^, and remain quenched when intercalated in (CGA)7/(TCG)7 and loaded in SPN. It suggests that release may not occur via simple diffusion, but rather by involving unwinding or enzymatic degradation of the DNA duplex. The slightly higher Dox accumulation in cells treated with SPN/siRNA/Dox compared with free Dox may reflect active export mechanisms for free Dox in cancer cells.

Apoptosis is a programmed cell death process and plays a critical role in maintaining homeostasis in normal tissues^33^. Most of the chemotherapeutics induce cell death via apoptosis. However, chemotherapeutics are often associated with resistance that alters the apoptotic pathways in cancer cells^34^. RRM2 plays important roles in DNA repair and gene replication, and its overexpression is linked to cell malignancy and resistance to DNA-damaging drugs such as tamoxifen and Dox^35,36^. Therefore, we investigated RRM2 downregulation combined with a DNA-damaging drug as a synergetic anticancer strategy. Apoptosis assays in MDA-MB-231 cells demonstrated a pro-apoptotic effect of SPN/siRRM2 compared with free Dox. Furthermore, cells treated with SPN/siRRM2/Dox exhibited a synergistic effect with the highest apoptosis, indicating that co-delivery overcomes Dox-induced drug resistance. Consistent with apoptosis data, SPN/siRRM2/Dox also resulted in increased cytotoxicity, further verifying the synergy between siRNA-mediated RRM2 silencing and chemotherapeutic treatment.

Importantly, Dox loading and release can be tuned by modifying the length of the DNA duplex or designing DNA nanostructures with higher CG content. This co-delivery strategy could also be further extended to other delivery carriers. Further studies combining this co-delivery strategy with cancer-targeting vesicles may further enhance selectivity and anti-cancer efficacy.

## Conclusions

In summary, we developed a simple and effective co-delivery system using a single-step co-encapsulation of siRNA and Dox-intercalated oligodeoxynucleotides with the cationic niosome. This system demonstrated several advantages: (1) Controllable loading of Dox and siRNA for potential personalized therapy; (2) efficient co-delivery of both Dox and siRNA into cancer cells; and (3) enhanced inhibition of cancer cell proliferation through synergistic effect. These findings highlight the potential of combining chemotherapeutics with antiproliferative siRNA for improved cancer treatment and suggest that this co-delivery strategy serve as a promising platform for future anticancer therapies.

## Conflict of interest

The authors declare no conflicts of interest.

## Acknowledgements

This work was supported by the Lundbeck Foundation (Project No. 23750). We additionally thank the Novo Nordisk Foundation (NNF) for their support through the RNA-Meta Center (NNF23OC0081177) and the CEMBID program (NNF17OC0028070). The authors also thank Anne F. Nielsen for helpful comments on the manuscript. We also acknowledge the China Scholarship Council (CSC) PhD scholarship from the Ministry of Education of China.

---

**Scheme 1.**
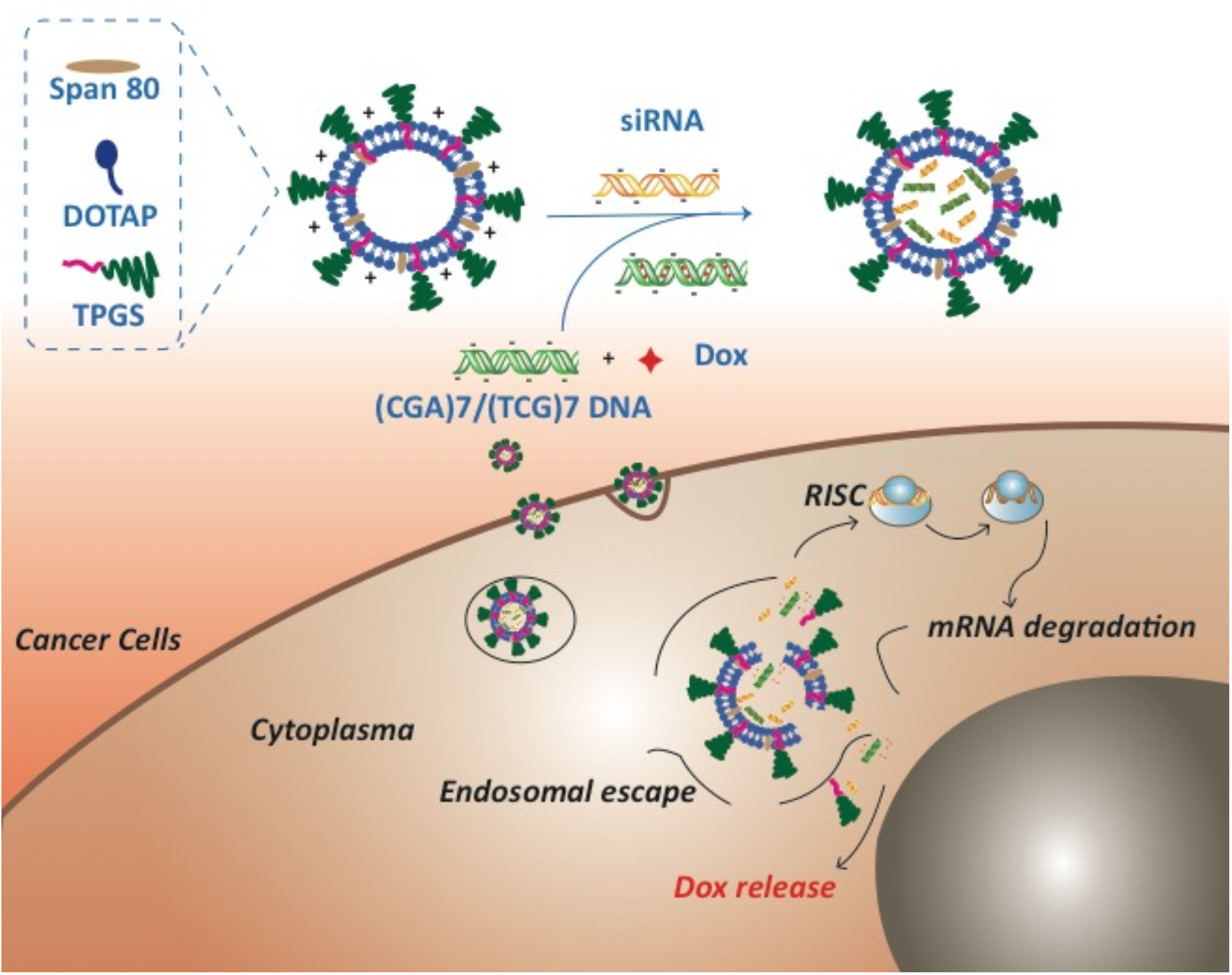
Schematic illustration the synthesis of co-delivery system (SPN/siRNA/Dox) by one-step encapsulating of siRNA and Dox and the endocytosis of SPN/siRNA/Dox and intracellular functionalization in cancer cells.

## Notes

### Competing Interest Statement

The authors have declared no competing interest.

