## supporting information for "Highly controllable co-delivery of siRNA and doxorubicin via cationic niosomes for synergistic anticancer effects"

**Electronic Supplementary Material (ESI)**

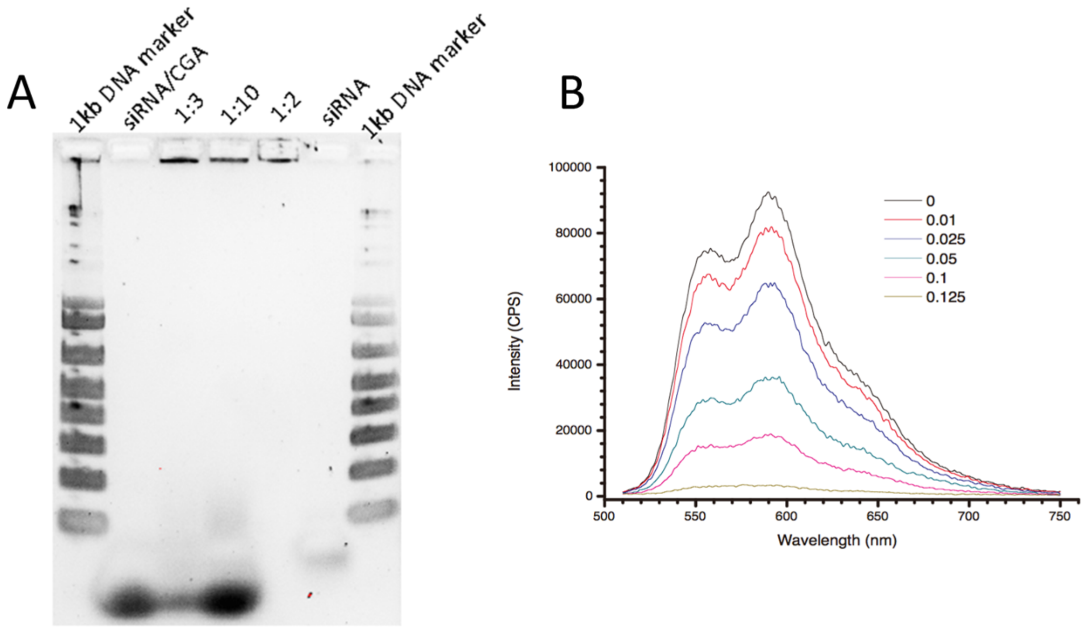


Fig. S1. (A) Native PAGE gel electrophoresis of SPN particles encapsulated with siRNA and CGA-Dox at various of weight ratios (volume ration siRNA:SPN):X, X and a X. Gel was stained with SYBR Gold. (B) Fluorescent emission spectrum of CGA-Dox at different molar ratios (Dox : CGA from 0 to 0.125).
